# Measles vaccine virus RNA in children more than 100 days after vaccination Running title: Measles vaccine virus RNA 100 days after vaccine

**DOI:** 10.1101/548701

**Authors:** Jamie McMahon, Ian M Mackay, Stephen B Lambert

## Abstract

Measles vaccines have been in use since the 1960s with excellent safety and effectiveness profiles. Limited data are available on detection of measles vaccine virus (MeVV) RNA in human subjects following vaccination. Available evidence suggests MeVV RNA can be identified up to 14 days after vaccination, with detection beyond this rare. In routine diagnostic testing, we used two real-time reverse transcription-polymerase chain reaction (RT-rPCR) assays for identifying measles virus (MeV) and MeVV RNA, followed by sequence confirmation of RT-rPCR positives by a semi-nested conventional RT-PCR. We report detection and confirmation of MeVV RNA from the respiratory tract of 11 children between 100 and 800 days after most recent receipt of measles-containing vaccine. These novel preliminary findings emphasize the importance of genotyping all MeV detections and highlight the need for further work to assess whether persistent MeVV RNA represents viable virus and if transmission to close contacts can occur.

## Introduction

Measles virus (MeV) is one of the most infectious pathogens of humans. It is highly transmissible with 90% of non-immune contacts exposed becoming infected; a high level of population immunity is required to interrupt transmission [1, 2]. MeV typically causes an unpleasant, self-limited febrile illness, however, complications can occur in up to 30% of cases, and in rare cases, can result in death [3].

Live attenuated measles vaccines have excellent safety and effectiveness profiles [2]. Use of measles vaccine has contributed to substantial declines in child mortality and morbidity: its use between 2000–2016 prevented an estimated 20.4 million deaths globally, a reduction of 84% in measles-associated deaths worldwide over the period [4].

Measles virus epidemiology is monitored using molecular typing, with eight clades designated A–H containing 24 recognised genotypes [5]. Measles vaccines were developed in the 1960s from genotype A wildtype viruses. Following global vaccine use, wild genotype A is now considered inactive with genotype A detections exclusively vaccine-associated [5].

Measles has been well controlled in Australia for more than a decade, with elimination of endemic measles transmission likely to have occurred as early as the turn of the century [6]. In March 2014, Australia was certified by the World Health Organization (WHO) as having eliminated measles [7].

Information about the detection of measles vaccine virus (MeVV) in human clinical samples is limited. Here we report a series of MeVV RNA detections, identified by testing specimens submitted for diagnostic measles PCR, more than 100 days following vaccination in Queensland, Australia.

## Methods

Our publicly-funded virology laboratory (Forensic and Scientific Services) receives requests for routine diagnostic measles PCR directly from primary care physicians and hospitals, or via private pathology providers, in Queensland and northern New South Wales, Australia. Between 2013 and 2017, our laboratory was the only laboratory in Queensland routinely offering measles PCR testing. Detailed illness and medical history from tested individuals were not routinely available. The work reported in this paper was deemed to be a quality assurance activity as described in the National Statement on Ethical Conduct in Human Research and was granted a waiver of ethics review by the Children’s Health Queensland Human Research Ethics Committee [8]. The Forensic and Scientific Services Human Ethics Committee assessed this project as not requiring full HREC review, as it was not recognised to be research, and is an audit of practice in accordance with the definition of research in the National Statement on Ethical Conduct in Human Research (2007) updated in 2018 [8].

Upon receipt of specimens with a request for MeV PCR, total RNA was extracted using the Qiagen EZ1 Mini kit v 2.0 (Qiagen, Hilden, Germany). Five of the 90μl elution was added to a measles virus (MeV) real-time reverse transcription-polymerase chain reaction (RT-rPCR) targeting the fusion (F) gene and designed to detect all MeV genotypes [9]. Nucleic acids were re-extracted from all MeV positive samples and testing was repeated using the MeV RT-rPCR. Additionally, the extracts from positive samples of patients with no known measles contact history, recent vaccination history, or at the request of the local public health unit, were further tested with a measles vaccine virus (MeVV) RT-rPCR targeting the matrix protein (M) gene designed specifically to detect MeVV strain, genotype A.

The primer, probes, and cycling conditions for the MeV and the MeVV RT-rPCR assays have been previously described [9]. The MeV RT-rPCR was used with a final primer concentration of 0.3μM and probe concentration of 0.15μM. Primers and probe were used at a final concentration of 0.3μM in the MeVV RT-rPCR. Samples were reverse transcribed for 5 min at 50°C (SuperScript^®^ III Platinum^®^ One-Step qRT-PCR Kit, Invitrogen, Carlsbad, CA), incubated at 95°C for 2 min, and then amplified using 40 cycles of 95°C for 3 s and 60°C for 30 s using either the Rotor-Gene Q or Rotor-Gene 6000 platform (Qiagen, Hilden, Germany).

Synthetic probe and primer controls were used for both RT-rPCR assays, alongside no template controls (NTC) and negative extraction controls [9]. A threshold cycle (C_T_) value of ≤40 indicated detection of MeV RNA whereas C_T_ values >40 were used to define a negative result.

To confirm RT-rPCR positive samples we used a semi-nested conventional RT-PCR requiring the generation of a 450-nucleotide (nt) fragment encoding the C-terminus of the nucleoprotein (N) gene [9]. First-round RT-PCR amplification was performed after adding 5μl of RNA extract to SuperScript™ III One-Step RT-PCR System with Platinum™ Taq DNA Polymerase (Invitrogen, Carlsbad, CA) mixes. A second round of amplification was performed by transferring 5μl of 1:100 diluted Round 1 product into 15μl Bioline MyFi™ Mix (BIOLINE GmbH, Luckenwalde, Germany) reactions.

Initial N-gene amplicon analysis was performed with 0.2 μl Round 2 product using the QIAxcel (Qiagen, Hilden, Germany). For positive reactions, gel electrophoresis of the remaining amplicon was followed by purification of the excised band using QIAquick Gel Extraction Kit (Qiagen, Hilden, Germany). Dye terminator sequencing was performed on the CEQ8000 Genetic Analysis System (SCIEX, Framingham, MA) using the GenomeLab DTCS Quick Start Kit (SCIEX, Framingham, MA).

Raw sequence data were cleaned, forward and reverse strands were aligned, and primer sequences removed. Sequences were aligned with the WHO designated MeV reference strains and phylogenetic analysis was conducted using Geneious version 11.1. A phylogenetic tree was constructed with MEGA7 using neighbor-joining method and maximum likelihood method using a bootstrap analysis of 1000 replicates. Molecular characterisation of the samples was confirmed using the MeaNS database genotyping tool [5].

Information on other virus PCR testing performed by us or other diagnostic laboratories who initially received the specimens (either the same specimen or specimens collected around the time of the positive MeVV specimen) was collated (Table 1).

**Table 1:** Results from other PCR testing performed on respiratory specimens in referring laboratories using the same specimen or a specimen collected around the time of the positive measles specimen.

| Case | PCR testing requested | Detections |
| --- | --- | --- |
| 1 | RSV, IFAV, IFBV, HPIV-1, HPIV-2, HPIV-3, HMPV, RV, AdV | HMPV |
| 2 | Not tested for other viruses | — |
| 3 | RSV, IFAV, IFBV, HPIV-1, HPIV-2, HPIV-3, HMPV, RV, AdV | RSV |
| 4 | RSV, IFAV, IFBV, HPIV-1, HPIV-2, HPIV-3, HMPV, RV, AdV | AdV |
| 5 | RSV, IFAV, IFBV, HPIV-1, HPIV-2, HPIV-3, HMPV, RV, AdV | RSV, HPIV-3 |
| 6 | RSV, IFAV, IFBV, HPIV-1, HPIV-2, HPIV-3, HMPV, RV, AdV | ND |
| 7 | Not tested for other viruses | — |
| 8 | RUBV*, RSV, IFAV, IFBV, HPIV-1, HPIV-2, HPIV-3, HMPV, RV, AdV | ND |
| 9 | RUBV* | ND |
| 10 | RSV, IFAV, IFBV, HPIV-1, HPIV-2, HPIV-3, HMPV, RV, AdV | AdV |
| 11 | Not tested for other viruses | — |

AdV = adenovirus; HPIV-1 = human parainfluenza virus type 1; HPIV-2 = human parainfluenza virus type 2; HPIV-3 = human parainfluenza virus type 3; HMPV = human metapneumovirus; IFAV = influenza A virus; IFBV = influenza B virus; ND = other viruses not detected; RSV = respiratory syncytial virus; , RUBV = rubella virus; RV = rhinovirus;
\* rubella virus PCR testing performed in our laboratory

## Results

Between 2013–2017, 9,940 samples were received for routine diagnostic MeV PCR from Queensland and northern New South Wales. During this period MeVV RNA was detected in 141 samples.

In 2014, a local public health unit initiated a follow-up process for a patient who was MeV RT-rPCR positive and had no risk factors for measles infection, including no recent travel or contact history. The patient had presented with a mild clinical illness that was not indicative of MeV infection. Further testing with the MeVV RT-rPCR followed by sequencing confirmed MeVV RNA was present in this sample collected 548 days after most recent measles vaccination.

Having identified an unusual positive case, we sought to identify and characterize further genotype A positives in submitted specimens. Between 2013–2017, MeVV strain RNA was detected in specimens from 11 children aged 18 to 45 months, more than 100 days (range: 101–784 days) following receipt of the most recent measles-containing vaccine (Table 2). Vaccines received included M-M-R II (Merck Sharp & Dohme Corp, West Point, Pennsylvania, US) (4 children), Priorix (GSK Biologicals, Rixensart, Belgium) (5 children), and Priorix-tetra (GSK Biologicals, Rixensart, Belgium) (2 children).

**Table 2.**
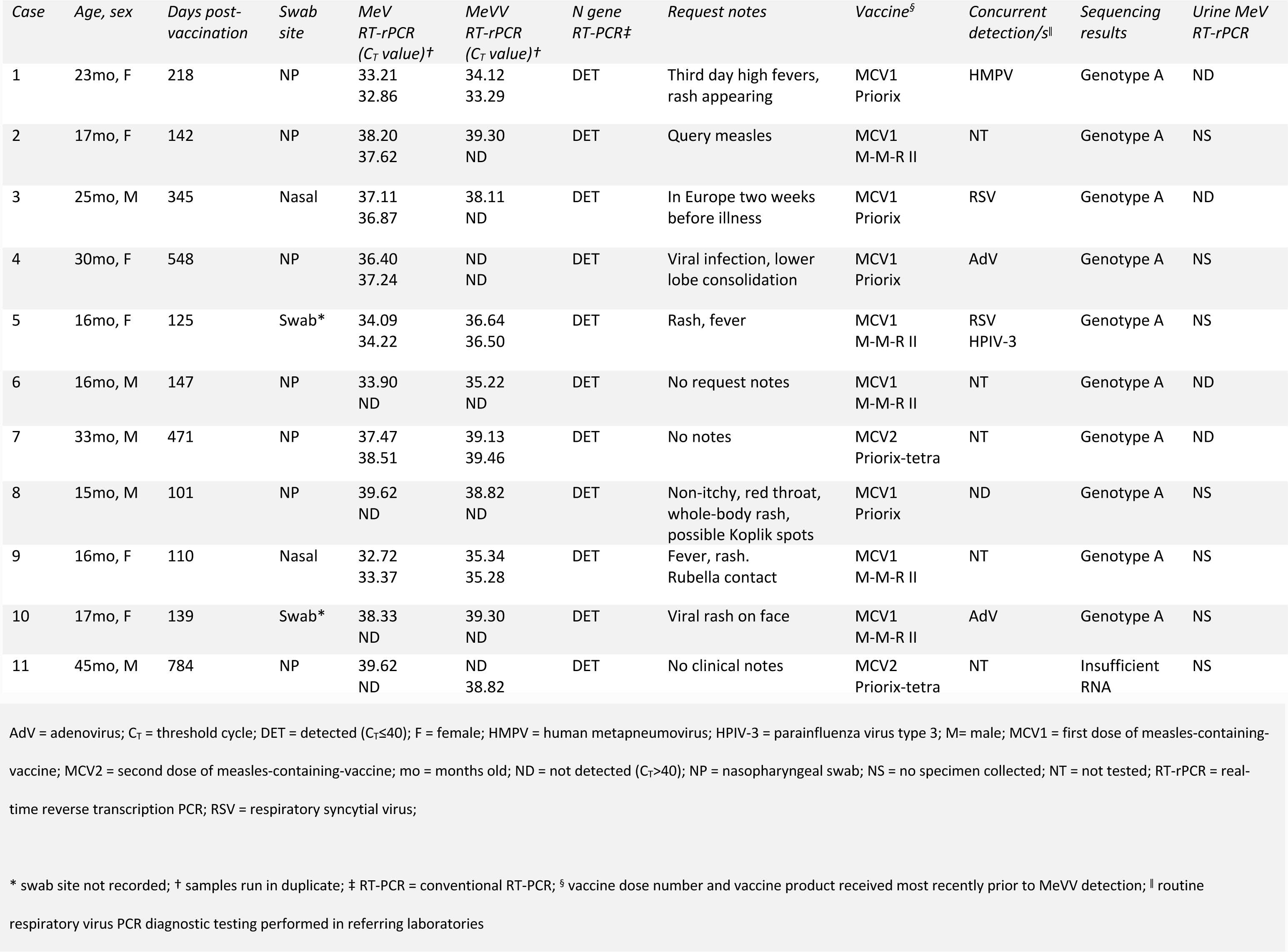
Details of cases with measles vaccine virus detection more than 100 days post-vaccination.

All 11 cases were initially detected using the MeV RT-rPCR. Ten cases were positive using the MeVV RT-rPCR. Case 4 was not detected using the MeVV RT-rPCR, however MeVV was confirmed by sequence analysis which identified MeV genotype A. We further confirmed MeV genotype A by sequencing in 10 cases (Figure 1). The remaining case, case 11, was initially not confirmed using the N gene RT-PCR. Upon repeat testing, a faint DNA fragment was present but could not be sequenced. The weak signal was likely due to the low level of RNA present in this extract (F gene RT-rPCR C_T_ value: 39).

**Figure 1:**
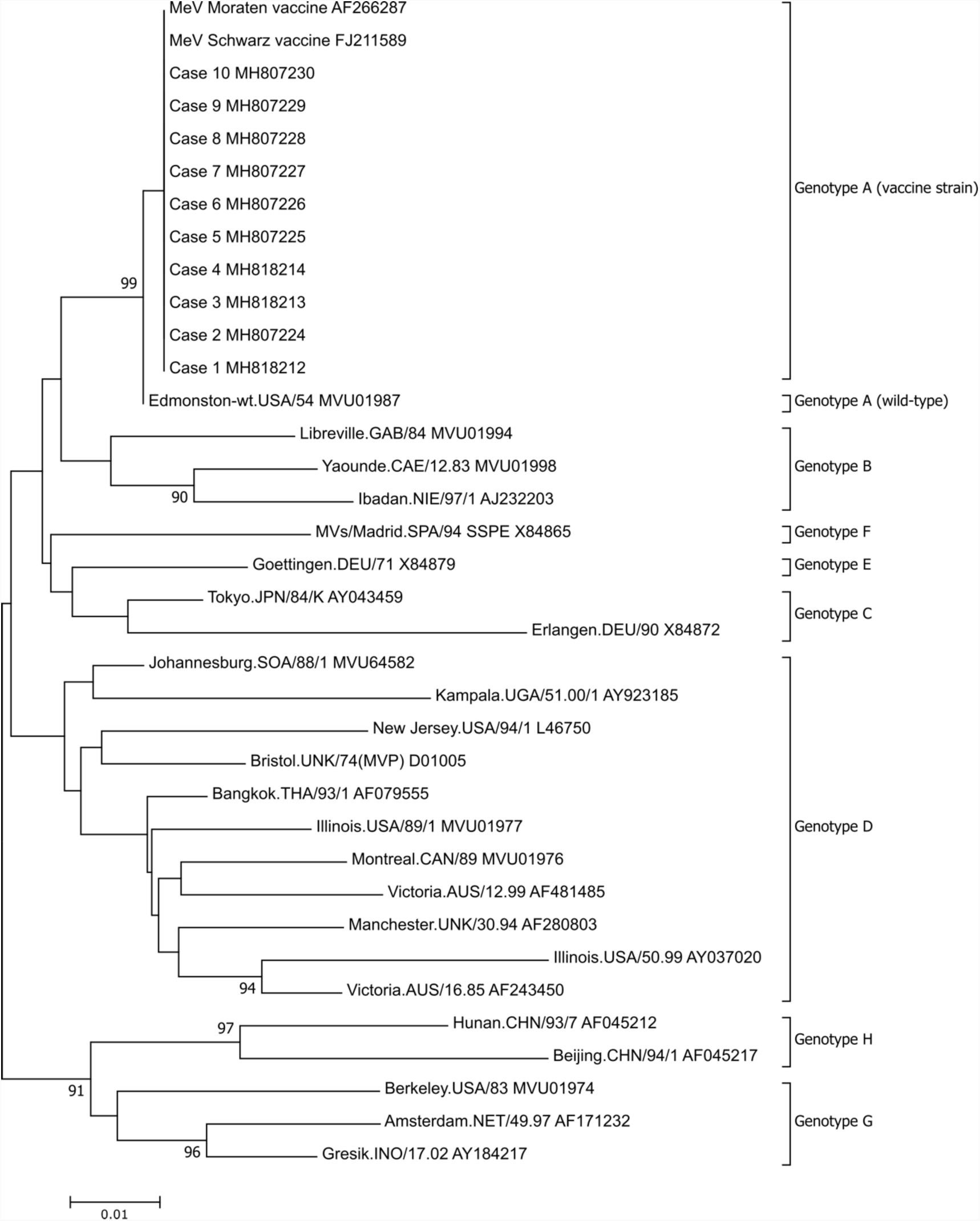
Phylogenetic analysis of MeV partial nucleoprotein (N)-gene sequences.

MeVV RNA was detected from nasopharyngeal (7), nasal (2), and unspecified respiratory swab sites (2). Five children had concurrent non-MeV respiratory virus detections; respiratory syncytial virus (1), respiratory syncytial virus and human parainfluenza virus 3 (1), adenovirus (2), and human metapneumovirus (1) (Table 1).

Of the 11 patients, four had urine collected concurrently. MeV RNA could not be detected in any of these samples. Case 9, who had MeVV RNA detected 110 days post-vaccination had both a throat and a nasal swab collected. The nasal swab was positive, but the throat swab was not. Ten cases confirmed by sequence analysis were determined to be genotype A (Figure 1).

The sequenced region is the World Health Organization (WHO)-recommended 450nt nucleoprotein (N) gene fragment encoding the carboxyl-terminal of the gene. WHO MeV reference strain sequences include GenBank accession numbers. MeVV sequences associated with patients are indicated by case number and associated GenBank accession number. Phylogenetic analysis used the Neighbor-Joining method Maximum Composite Likelihood methods was performed in MEGA7 with a bootstrap analysis of 1000 replicates. The percentage of replicate trees in the bootstrap test (1000 replicates) greater than 85 are shown next to the branches. A higher resolution version is located at https://doi.org/10.6084/m9.figshare.7649063.

## Discussion

During routine PCR screening in response to clinical requests for MeV testing, we found genotype A MeV RNA in submitted respiratory specimens between 100 and 800 days after most recent vaccination date from 11 children over a period of five years. We confirmed RNA detection using a MeVV-specific RT-rPCR assay, followed by use of a semi-nested conventional RT-PCR and nucleotide sequencing. Findings of this nature, in any human clinical sample type so long following documented vaccination, have not been described previously.

MeV detection has been infrequently described several months following infection and detected from a range of tissues. Persistent, often mutated, MeV infection causes rare, sometimes fatal, complications, including measles inclusion body encephalitis and subacute sclerosing panencephalitis many years post-infection [10]. MeV RNA, typing not further described, was detected in ileal biopsies from two children at least six months following receipt of their most recent measles-containing vaccine, but no respiratory sites were tested [11]. Delayed MeV detection has been reported in non-respiratory tissues: one paper describes the presence of MeV RNA in lymphocytes of healthy people who had a wildtype infection up to 30 years prior [12]. However, sequence analysis in this study suggested recent, asymptomatic infection of MeV rather than MeV persistance. MeV RNA has been detected in bone marrow aspirates taken for the diagnosis of malignant bone diseases [13]. MeV RNA has been found in autopsy tissue from apparently healthy people using PCR with detections in the brain, lung, liver, kidney, and spleen (8–20% detection prevalence) [14].

Data on MeVV detection in human clinical samples is limited: this may be because specific testing for MeVV is uncommon, detection of MeVV is uncommon, symptoms are not associated with presence of MeVV RNA, or a combination of these. Unlike wildtype MeV, MeVV sequences are not submitted routinely to the WHO Measles Nucleotide Surveillance (MeaNS) database [5]. MeVV detection up to 14 days after vaccination is reported [15-17]. There is one case report of detection from the nasopharynx five weeks post-vaccination [18]. Vaccine-associated genotype A has also been previously detected in urine 25 days following vaccination [19]. Our description of this small case series is likely made possible for a number of reasons: enhanced surveillance involving increased requests for testing in an elimination setting; the application of increasingly sensitive PCR assays to many thousands of specimens; and the ability to rapidly detect and differentiate wildtype MeV from MeVV.

We report these preliminary findings as an initial communication, but recognise the limitations in our work. Firstly, we have reported data on only 11 episodes of MeVV RNA detection. From this it is not possible to determine if MeVV RNA may be intermittently detectable in some or most vaccinated individuals, or whether we are reporting a truly rare event. Also, due to small case numbers, it is unclear if these detections are specifically associated with concurrent presence of other respiratory viruses. Finally, detection of MeVV RNA alone does not allow for any assessment of whether infectious virus is present.

Further work is required to expand on our preliminary findings. This work includes assessing whether the MeVV RNA detected derives from viable virus, or an RNA transcription process that produces non-infectious or no virus. If live virus is produced it will be important to assess if transmission to close contacts occurs. It is worth noting there is no documented evidence of human-to-human transmission of MeVV [20]. These further details are required to understand the significance of our findings, and their implications for immune boosting, disease control, and reaching and maintaining WHO measles elimination targets. Our findings also highlight the importance of genotyping all MeV detections, not only to provide laboratory evidence of the absence of endemic MeV circulation, but to document the prevalence and detailed epidemiology of genotype A detection.

## Funding

This work was supported by Forensic and Scientific Services, Health Support Queensland. This research did not receive any specific grant from funding agencies in the public, commercial, or not-for-profit sectors.

## Acknowledgements

We thank Public Health Virology members Frederick Moore, Glen Hewitson, Doris Genge, Judy Northill, Amanda De Jong, and Peter Burtonclay who contributed to the laboratory confirmation of the cases. We also thank the staff in the public health units involved in following up these cases, and the families of the cases.

## Conflicts of Interest

Jamie McMahon – No conflict

Ian M Mackay – No conflict

Stephen B Lambert – SBL is the current Chair of the national Measles and Rubella Elimination Working Party

## Non-standard abbreviations used in the article

MeVV: measles vaccine virus
MeV: measles virus

